# 3D Imaging of Honeybee Swarm Assembly and Disassembly

**DOI:** 10.64898/2026.03.17.711698

**Authors:** Danielle L. Chase, Daniel Zhu, Mahi Kathait, Henry Robertson, Jash Shah, Sully Harrer, Gary Nave, Nolan R. Bonnie, Orit Peleg

## Abstract

When honeybee colonies reproduce by fission, several thousand bees and their queen depart the parental nest and temporarily form a dense cluster on a tree branch or other surface while searching for a new nest site. Once the new nest site is selected, the swarm disassembles and flies toward it. How honeybees transition rapidly between dispersed flight and an aggregated cluster remains an open question. Here, we develop an experimental system and three-dimensional imaging pipeline to track individual flying bees together with the evolving morphology of the swarm during formation and dissolution. We report results from a representative swarming event. During assembly, swarms rapidly form low-density clusters before undergoing a slower contraction to a more dense steady state configuration. In contrast, disassembly occurs significantly faster than assembly and is characterized by strongly divergent flight, with bees departing the swarm in all directions. Overall, this method is able to demonstrate the coupled flight and morphological dynamics that underlie honeybee swarm assembly. Because the system is relatively low-cost and low-power, it is readily adaptable for three-dimensional imaging of other biological collectives in naturalistic environments.

## Introduction

Animal groups are capable of functions that individuals cannot achieve alone [17, 7]. Maintaining cohesiveness is essential for realizing the benefits of group living: when faced with decisions such as whether to stay or go, and where to go, groups must first reach consensus [4, 9]. Subsequently, coherent movement ensures that individuals remain together [10, 11, 2].

In dense biological aggregations, cohesion can arise through direct physical interactions between individuals [1, 35]. Army ants build bridges [30], bacteria form biofilms [8, 32], *Dictyostelium discoideum* form fruiting bodies [5], fire ants assemble into floating rafts [19, 16], and California blackworms form tangles [23, 20, 24]. These collective structures can reach sizes orders of magnitude larger than any of their constituent individuals and enable diverse functionalities ranging from locomotion and thermoregulation to reproduction. At the same time, many biological processes require the rapid dissolution of these assemblies. Thus, aggregations must have the capacity to maintain cohesion while also retaining the capacity to assemble and disassemble quickly.

Honeybee swarms are one such biological aggregation that forms a transient, dense collective. When a colony reproduces by fission, several thousand bees and their queen take flight from their nest and coalesce into a hanging cluster (Fig. 1A). This cluster serves as a temporary home for several hours to a day while scout bees search for a permanent nest site [18, 34, 33]. During this period, the swarm must buffer environmental perturbations like wind, rain, and temperature changes. The cluster adapts its density to thermoregulate [13, 12, 26] and maintains mechanical stability by tuning the spatial distribution of bees within the swarm [36] and adjusting its morphology in response to mechanical perturbations [25]. Once a nest-site consensus is reached, the cluster rapidly disassembles and takes flight as a coherent group [3] (Fig. 1E).

**Figure 1.**
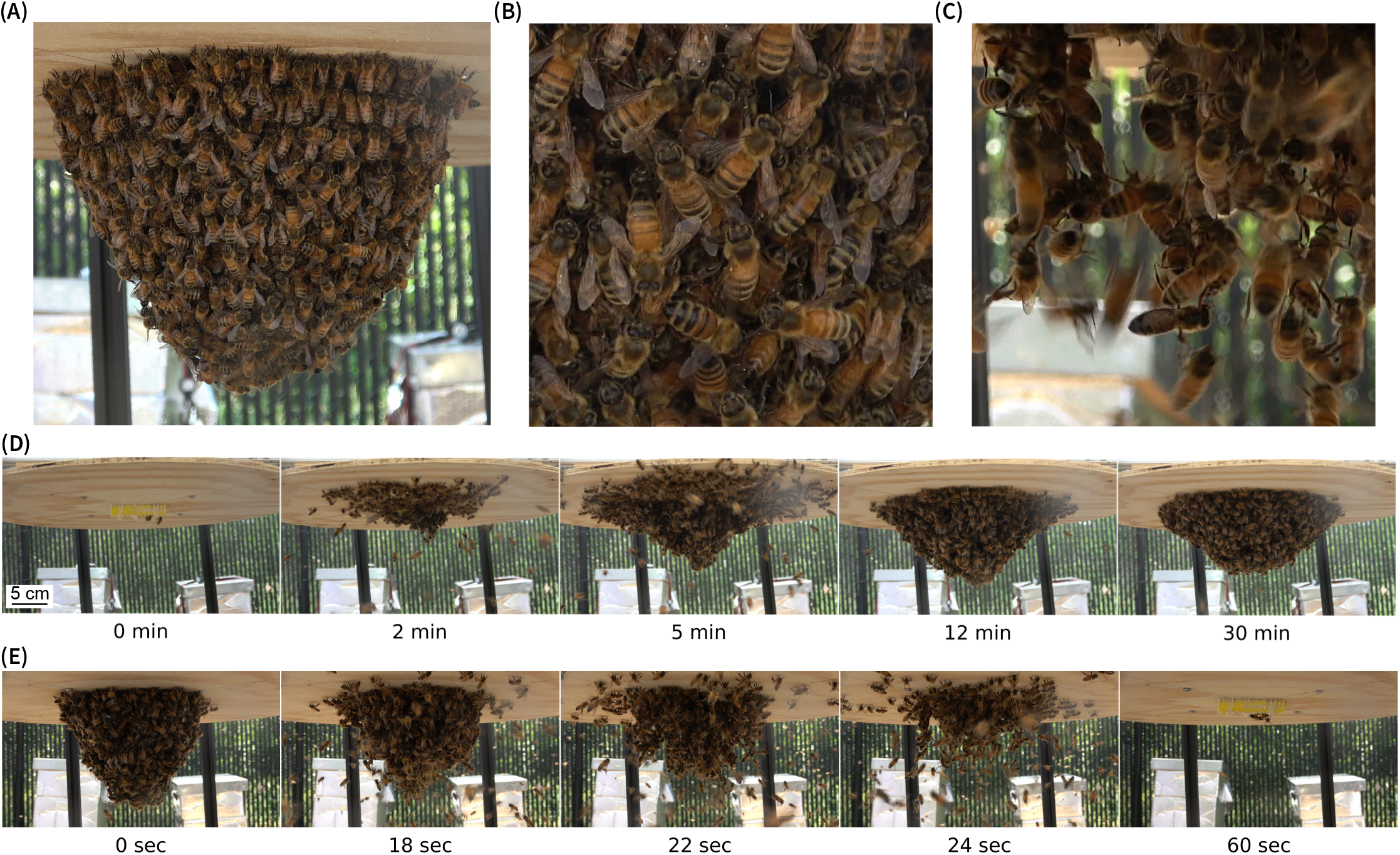
Honeybee swarm structure and dynamics. (A) A honeybee swarm hanging from the underside of a horizontal board in the characteristic pendant shape. (B) Close-up view of the surface of a stable swarm. (C) Close-up view of the swarm during disassembly, showing the connections between individual bees. (D) Time series illustrating the self-assembly of a swarm around a caged queen over 30 minutes. (E) Time series images of swarm disassembly over 60 seconds.

Despite being readily observable in nature, the dynamic processes underlying swarm assembly and disassembly are not well understood. Studying these transitions in the field poses several challenges. The onset of swarming depends on multiple biological and environmental factors, making its precise timing difficult to predict [28, 27], and swarm landing sites are often inaccessible, such as on high tree branches. In this work, we present an experimental study of swarm self-assembly in which we control the spatial and temporal components of swarming with a caged queen. Using stereoscopic imaging, we track the trajectories of individual bees and quantify the evolving swarm morphology. This approach enables quantitative investigation of a key question: How do swarms assemble and disassemble rapidly while maintaining mechanical stability?

## Methods

### Swarm preparation and maintenance

We use artificial swarms from colonies of European honeybees (*Apis mellifera* L.). A swarm is prepared by shaking *∼* 600 grams of bees (approximately 5,500 bees) from frames in a hive into a ventilated bucket. A caged queen from a queen bank is placed at the top of the bucket. The swarm is kept in a protected environment indoors for two days and fed *ad libitum* with 2:1 (volume/volume) sucrose solution delivered via a gravity feeder. This treatment induces behavior in the artificial swarm that is similar to a natural reproductive honeybee swarm [13, 33].

To assemble the swarm outdoors, the caged queen is attached to the underside of a horizontal wooden board, after which, the bucket is opened to allow the bees to cluster around her. During experiments, the swarm is fed at least twice daily by misting with a 1:1 sucrose solution. Bees also have unrestricted access to forage naturally during this time.

### Experimental setup

The swarm is placed under a tent with two side walls to provide shade and protection from the elements (Fig. 2A). To keep track of the number of bees in the swarm, the swarm hangs from the underside of a 50 cm diameter wooden board, which is suspended from a scale. The mass of the swarm is continuously monitored by a camera (GoPro MAX Action Camera) that records an image of the scale at 1 frame per second. To obtain stereoscopic views of the swarm, two pairs of GoPro MAX cameras are used: one pair positioned on one side of the swarm and an identical pair on the opposite side (Fig. 2B). In each pair, the optical axes are parallel and the cameras are separated by 50 cm and mounted to a tripod. The rig was positioned 80 cm from the front of the board and 105 cm from the center. Video is recorded from each camera continuously at 60 frames per second with a resolution of 1920 *×* 1440 pixels. Before each experiment, a video is recorded of a camera calibration board and a flashing LED light is recorded for frame synchronization.

**Figure 2.**
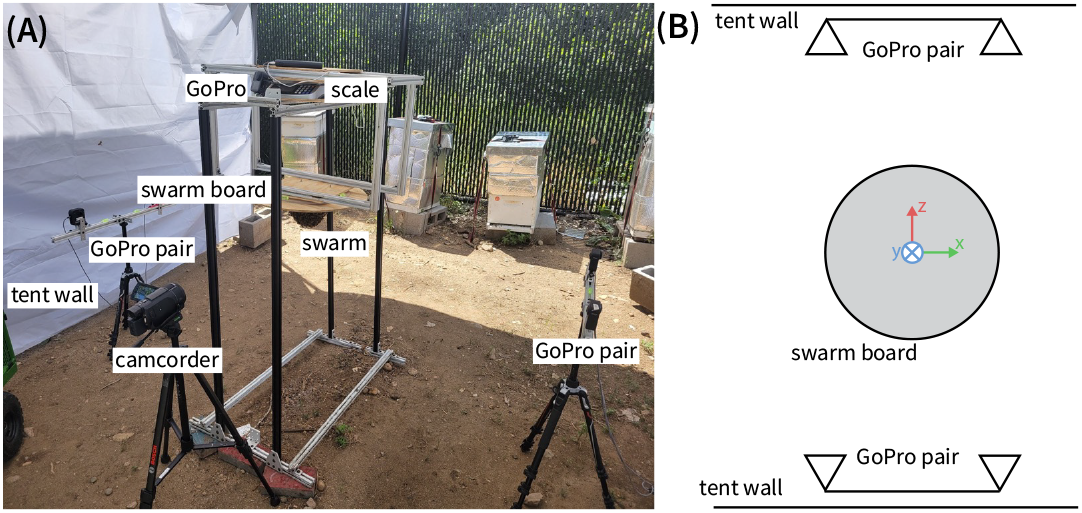
Experimental setup. (A) Photograph of the outdoor enclosure. A honeybee swarm hangs from the underside of a wooden disc, which is suspended from a scale to continuously monitor swarm mass. Two pairs of GoPro cameras positioned on opposite sides of the swarm record stereo video of the swarm, while a camcorder provides a higher resolution close-up view. (B) A schematic diagram showing the relative positions of the cameras and a coordinate system centered on the swarm board.

Video is acquired from approximately 8 AM to 6 PM daily to capture the swarming event in which the swarm cluster takes off and begins to fly to the selected nest site but, upon detecting the queen’s absence, returns and re-assembles into a cohesive cluster. This process typically occurs once per day if the weather conditions are good.

### Detection of individual bees and swarm morphology

While the white background makes it possible to do simple image difference to identify flying bees and the swarm boundary, we found that the machine learning models are significantly more robust against background fluctuations in lighting throughout a day and between days and to other changes in the environment.

The swarm boundary is segmented using the SAM2 model [29]. For each video frame, the model produced a binary mask of the swarm, and the outer contour of the mask is extracted to obtain the swarm boundary (Fig. 3A and C). Each video is segmented using positive and negative prompts on an initial frame and a few intermediate frames to inform the model of the object of interest.

**Figure 3.**
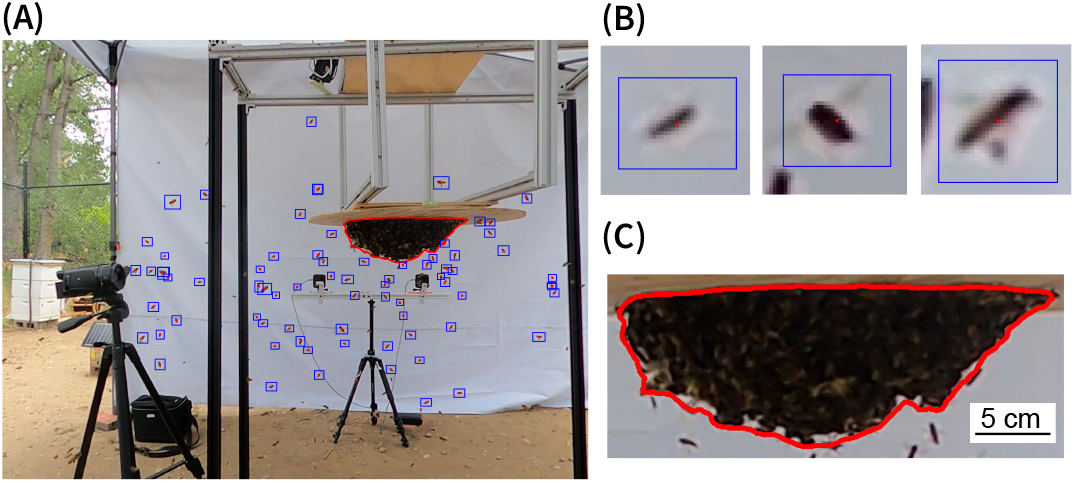
Detection of individual bees and swarm morphology. (A) Example video frame showing detections of flying bees (blue bounding boxes with red centroids) and the segmented swarm (red contour line). (B) Examples of individual bee detections. (C) Close-up view of the segmented swarm boundary used to extract swarm shape.

Flying bees are detected using a fine-tuned YOLO11 (Ultralytics) model [14]. The model was fine-tuned on labeled bee images using a 142:12 split for training and testing with images containing 10-100 bees. Fine-tuning was performed for 300 epochs using standard YOLO11 augmentations, including random flips, scaling, and HSV jitter. For each bee, the model predicts a bounding box whose centroid was then taken as the bee’s 2D position in the image (Fig. 3A and B).

### 3D reconstruction of bee trajectories

Honeybees approach and join the forming swarm cluster along three-dimensional trajectories that cannot be inferred from a single camera view. Depth information is required to resolve the full trajectories and velocities of flying bees as well as the position and evolution of the swarm cluster. Stereoscopic imaging overcomes these limitations by providing paired views that enable geometric triangulation of bee positions and the swarm boundary throughout assembly and disassembly. Stereoscopic approaches have been widely used for tracking the 3D trajectories of flying insect swarms [15, 37, 31].

A bee’s 3D position can be reconstructed by triangulating its corresponding locations in the left and right camera views. In a pair of calibrated cameras, each pixel location corresponds to a ray in 3D space, and the desired point lies at the intersection of the two rays. We work in rectified coordinates, so corresponding points lie along horizontal epipolar lines and the horizontal disparity between the left and right images determines the depth coordinate *Z*. In principle, triangulated 3D points can be linked together in time to form a trajectory. However, the high spatial density of flying bees makes linking 3D points into stable trajectories difficult. This motivates a two-step procedure: first constructing 2D trajectories from image coordinates in the left camera view, and then finding the corresponding right camera detections frame-by-frame to triangulate 3D positions (Fig. 4). In the detection of flying bees (Fig. 3), individuals do not maintain persistent identities between frames, so we link detections into trajectories with frame-by-frame tracking. First, we predict the current position of each active track *i* via 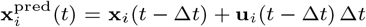, where the velocity **u**_*i*_(*t −* 1) is estimated using an exponential moving average of recent frame-to-frame velocity measurements for that track. If only a single detection exists, the last position is reused. Each detection in the current frame is **x**_*j*_ (*t*) = (*x*_*j*_, *y*_*j*_). For each predicted track position 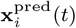, detections satisfying 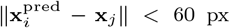. For these candidates, the assignment cost is defined as 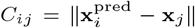. The full cost matrix **C** is constructed for all valid candidate pairs and the Hungarian algorithm is used to compute the minimum-cost assignment between left tracks and right detections. Unassigned detections initiate new trajectories, while tracks without a matching detection for two consecutive frames are terminated. The resulting 2D trajectories are stored as (*k, x, y, t*) for each track *k*.

**Figure 4.**
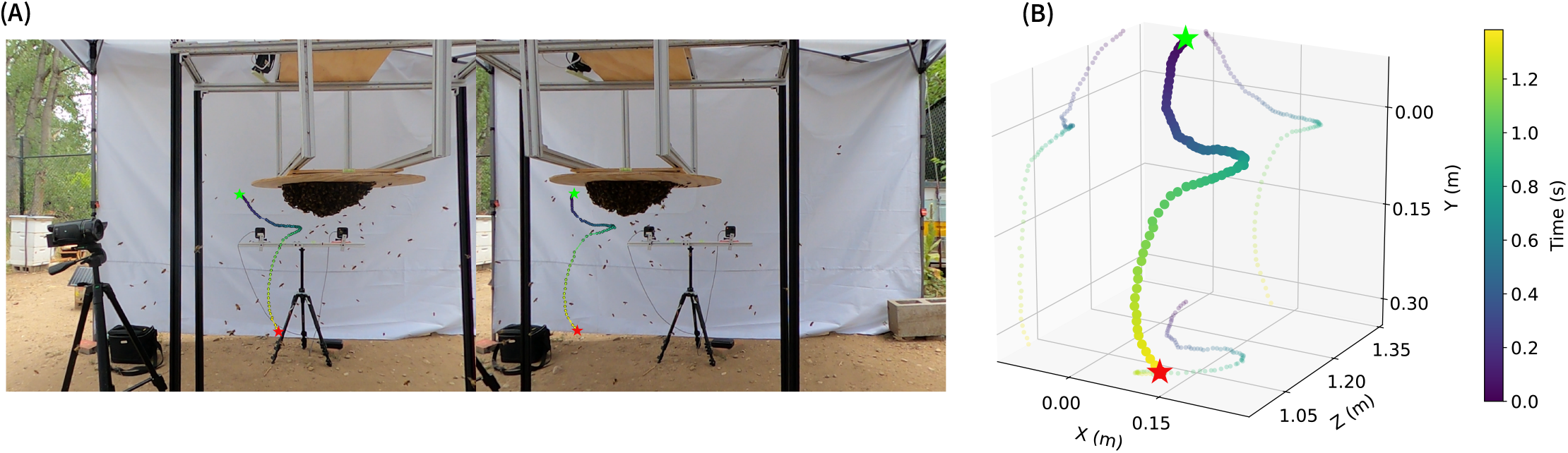
3D reconstruction of an individual bee flight trajectory. (A) Side-by-side frames from the left and right cameras of a stereo pair showing the tracked 2D positions of a flying bee. The start and end of the trajectory are indicated by the green and red markers, respectively. (B) Corresponding 3D trajectory reconstructed from stereo geometry, with color representing time.

To obtain 3D trajectories, every frame from the left camera supplies a set of active tracks, and the detections in the right camera supply the candidate matches (the camera choice for each is arbitrary). For each frame, we enumerate all candidate pairs between left-track positions (*k, x*_*L*_, *y*_*L*_) and right camera detections (*x*_*R*_, *y*_*R*_). We set a vertical threshold for possible matches of 6 px and require the disparity *d* = *x*_*L*_ *− x*_*R*_ to be within a range corresponding to a valid depth 0 *< Z <* 1.8 m. For each remaining candidate pair, we compute a cost representing its likelihood of being the true correspondence. For left tracks with an existing 3D state, we use a constant-velocity model to predict the track position, **X**^pred^(*t*) = **X**(*t −* Δ*t*) + **u**(*t −* Δ*t*)Δ*t*, where the velocity **u**(*t−*Δ*t*) is estimated using an exponential moving average of recent frame-to-frame velocity measurements. Allowable candidate matches must lie within 5 cm of the predicted position, and for these candidates, the assignment cost is defined as the Euclidean distance between the triangulated point and the predicted position, 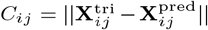. If the trajectory does not yet have a velocity prior, the cost is determined by the vertical epipolar residual *C*_*ij*_ = ∥*y*_*L*_−*y*_*R*_*∥*. We obtain left–right correspondences by solving the linear assignment problem defined by the pairwise matching costs for each frame. Accepted matches are triangulated into 3D points, which inherit the track identity from the left camera. For each resulting bee trajectory, we apply a 0.05 s (3-frame) moving average filter to reduce frame-to-frame noise in the detected bee positions and the resulting fluctuations in the triangulated depth.

### Measurement of Swarm Morphology

To quantify swarm morphology, we first obtain binary masks of the silhouette from the left and right stereo cameras (Fig. 3C). Although disparity can generally be computed wherever pixel-wise correspondences exist, in our images the interior of the silhouette is visually noisy, and the only matchable features are the boundary pixels. Accordingly, we compute disparity by matching boundary pixels along each image row. To approximate the three-dimensional position of the silhouette in each frame, we compute the mean depth of the boundary pixels and use this value to define the analysis plane for the 2D contour extracted from the mask. From this contour, we measure swarm diameter (maximum horizontal extent), swarm height (vertical extent), and projected area. To estimate the volume of the swarm, we assume that the cluster is approximately axisymmetric about its vertical axis and compute the volumetric estimate obtained by revolving the 2D silhouette around this axis.

## Results and Discussion

We quantify the temporal evolution of swarm morphology and flight activity during self-assembly and disassembly (Fig. 5A–D). To characterize flight dynamics, we count the number of visually detected flying bees in each frame (purple curve in Fig. 5A). This measure does not represent a full count of all flying bees since some bees are out of frame, but rather a consistent measure of the bees flying nearby the swarm cluster. When the swarm is stable, very few bees are flying, only the occasional scout or forager leaving or returning to the swarm. At the onset of disassembly *t* = 0, the visually detected number of flying bees rises sharply, which is coupled with the rapid drop in swarm mass (red curve in Fig. 5A). Within approximately 60 seconds, the cluster is fully detached from the board and nearly 0 bees are left behind as the swarm takes off to the nest site.

**Figure 5.**
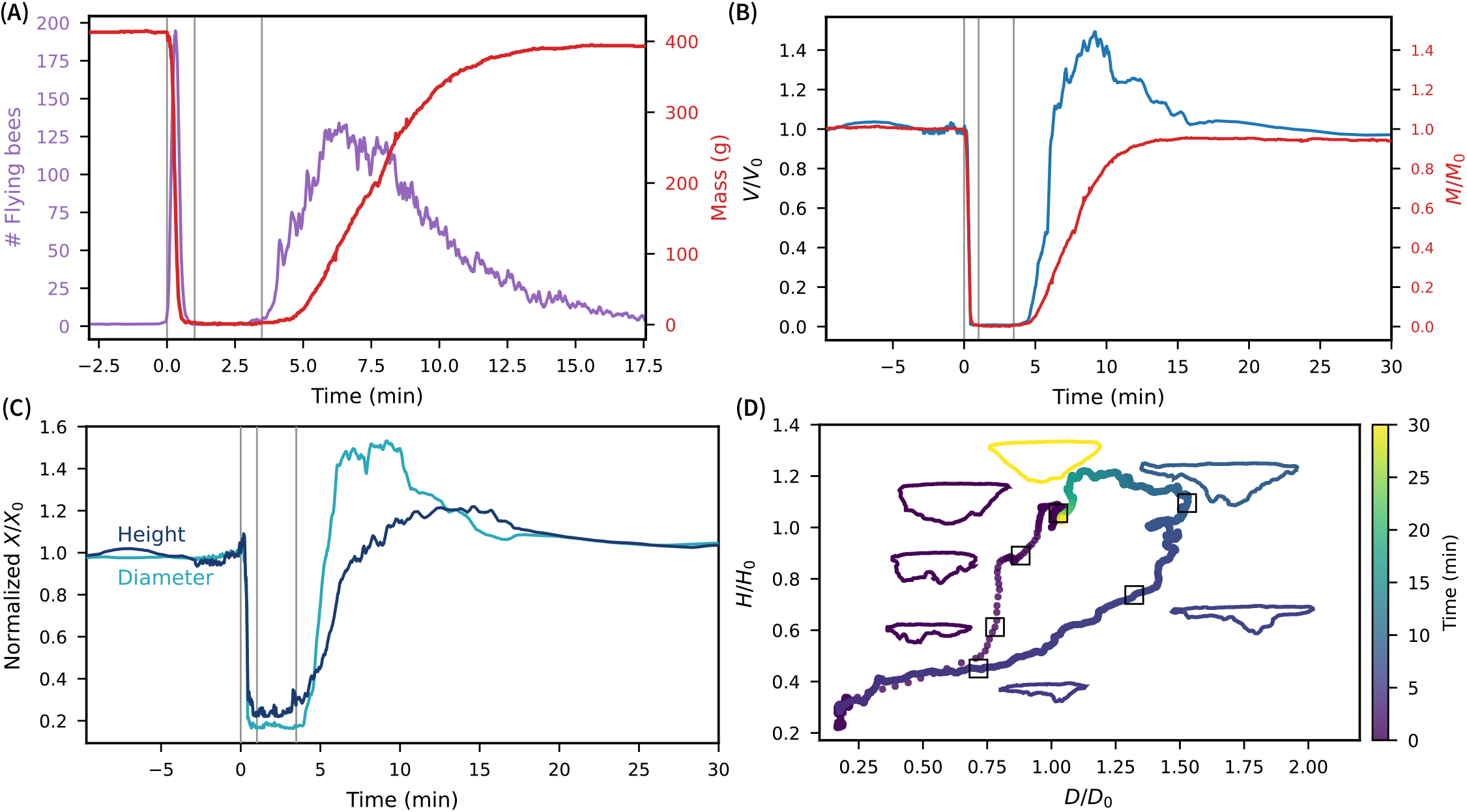
Dynamics of swarm morphology. (A) Number of detected flying bees (purple) and swarm mass (red) over time, showing the rapid disassembly followed by subsequent assembly. Vertical lines indicate the start of disassembly (time 0, defined as a mass decrease of three grams below the maximum), the end of disassembly (when no bees remain on the swarm), and the onset of assembly (defined as a mass increase of three grams above the minimum). (B) Normalized swarm volume *V/V*_0_ (blue) and mass *M/M*_0_ (red) during disassembly and assembly, with initial values measured at time 0. (C) Normalized swarm diameter *D/D*_0_ and height *H/H*_0_ over time. (D) Morphological trajectory during swarm self-assembly, with points colored by time and silhouettes indicating the swarm shape along the trajectory.

After a few minutes, likely upon realizing the queen is not with them, bees begin returning to the swarm location, initiating the assembly of the swarm around the caged queen. Overall, the formation process is much slower than the 60 second dissolution process. The early phase of aggregation is characterized by a high arrival rate of bees, and shortly after, a rapid increase in swarm mass as the flying bees attach to the board and the swarm cluster (Fig. 5A). Over time, the process slows, taking approximately 12 minutes to reach the final mass. Over the period of disassembly and subsequent reassembly, the swarm mass decreases by approximately 20 grams. This mass loss can be attributed to bees that do not return to the cluster (approximately 150 bees by mass) and to reductions in individual bee mass resulting from metabolic energy expenditure and water loss during flight. The experiments take place in an apiary with twenty other hives, the activity of bees from other colonies could create a visually and chemically cluttered environment that may make it more difficult for some bees to locate and return to the swarm site.

To capture morphological evolution, we measure the swarm volume *V* (*t*) and normalize it by the volume at the onset of disassembly *V*_0_ = *V* (*t* = 0) (Fig. 5B). During takeoff, the normalized volume *V/V*_0_ decreases rapidly as bees depart the cluster. To further characterize how the swarm shape evolves during this process, we compute the normalized diameter and height from the swarm silhouette (Fig. 5C). Prior to takeoff, both diameter and height fluctuate rapidly and exhibit a slight upward drift, potentially reflecting increased activity of bees on the surface of the swarm cluster. Once disassembly begins, both quantities decrease rapidly, although the height decreases faster than the diameter. The swarm height decreases by 50%, while the diameter decreases by only 25%. As a result, the cluster briefly adopts a wide, flattened shape in which bees remain primarily attached to the board before continuing to take off (Fig. 5D). The nonzero residual diameter and height correspond to the queen cage on the board (Fig. 1E).

During assembly, the volume initially increases rapidly, overshooting its initial value by 50%. This overshoot corresponds to a swarm that is less dense than the initial configuration at time 0. As the mass accumulation slows, the swarm begins to contract (Fig. 5B). After the swarm reaches its final mass, it continues to contract, not reaching its steady state value until approximately 25 minutes after assembly begins. During the initial volume increase, both the swarm height and diameter grow as bees aggregate (Fig. 5C). However, they increase at different rates: the diameter grows more quickly than the height, producing an initially broad, flat structure. The diameter overshoots its initial value by more than 50%, whereas the height overshoots by approximately 20%. The diameter then begins to contract, followed by the height, with both gradually relaxing back toward their initial configuration.

The morphological trajectory during assembly (Fig. 5D) reflects this progression. The swarm first grows by expanding its diameter rapidly while increasing in height more slowly. Subsequently, the height continues to increase while the diameter begins to contract, and finally, the height contracts. These morphological regimes reflect an underlying two-stage process in the bees’ local behavior. The first stage involves the rapid aggregation of bees and the second involves slower reorganization of bees within the swarm. Together with the volume overshoot, these trends suggest that the swarm prioritizes fast cohesion which could be advantageous given the energetic cost of flight. Once aggregated, the bees can prioritize a coherent and mechanically stable cluster by redistributing bees within the swarm. The faster increase in diameter relative to height may also reflect the stability of landing on the swarm board, which provides a large, flat surface for initial attachment before vertical extension occurs.

To relate the evolution of swarm morphology to the dynamics of individual bees, we measure the velocity field of flying bees in the vicinity of the swarm (Fig. 6A and C). From the 3D trajectories, we first compute instantaneous velocities, then bin the 3D space into 10 cm cubes and average the velocities of all trajectory points within each bin over the assembly and disassembly windows (6 minutes 49 seconds and 60 seconds, respectively). We also quantify the spatial density of trajectories by counting the number of unique trajectories passing through each 10 cm bin and depth-averaging the result (Fig. 6B and D). The resulting velocity and density fields reveal distinct patterns between the two phases.

**Figure 6.**
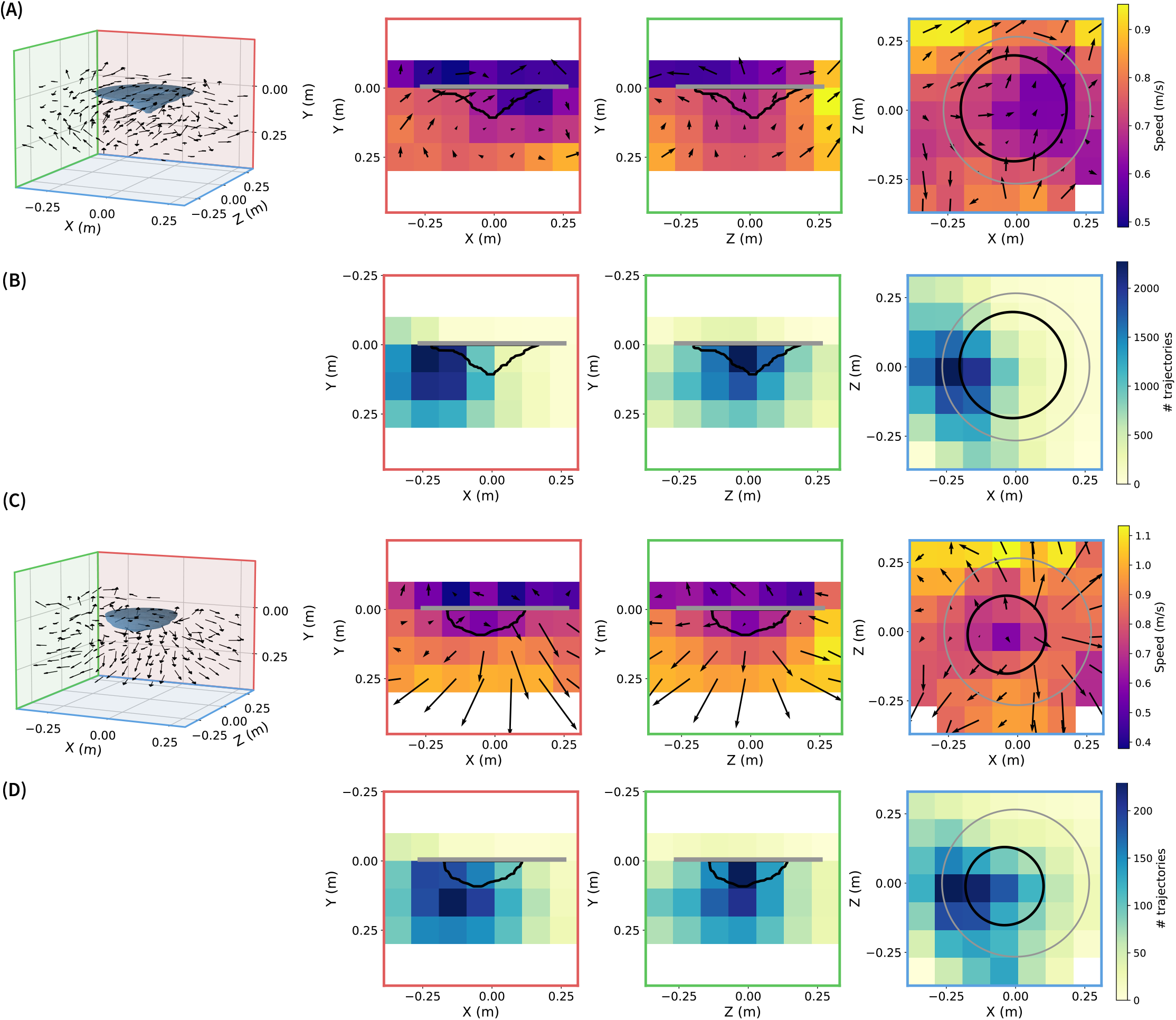
Velocity fields around the swarm during assembly and disassembly. (A) Reconstructed 3D velocity field of flying bees during swarm assembly, shown as the full 3D vector field (left) and in three orthogonal projections (right). The velocity field is computed from the bee trajectories collected over the first 6 minutes 49 seconds of assembly. The silhouette indicates the swarm shape at the end of this interval. In the orthogonal projections, data are averaged along the direction perpendicular to the projection plane. Color indicates the mean flight speed, and the length and direction of arrows show the mean in-plane velocity. The gray boundary indicates the swarm board, and the coordinate system origin is the position of the queen cage at center of the board. (B) Spatial density of flight trajectories reported as the number of unique trajectories that pass through each voxel during assembly. (C-D) Equivalent visualizations as (A-B) for the disassembly phase, over a period of 60 seconds (the full time period for disassembly). The silhouette shows the swarm shape at the start of disassembly.

During assembly (Fig. 6A-B), the velocity field is characterized by swirling motion around the swarm. The *XY* -projection shows net motion in the (+*X, −Y*) direction, suggesting that bees typically approach from below and fly upward toward the cluster. Bees that overshoot in the *X−*direction (*XZ*-projection) can recirculate to join the swarm. Bee density is highest in the *−X* half-space of the volume, where bees are approaching the swarm. As a result, accumulation is strongest on the *−X* side of the swarm, consistent with the early asymmetry in swarm shape (Fig. 5D).

During disassembly (Fig. 6C), the velocity field displays strong, outward-directed flows. Bees fly away from the swarm in all directions with relatively large magnitudes. This pattern reflects the rapid, coordinated takeoff of the cluster. Once the swarm decides to leave, thousands of bees depart along diverging trajectories. The approximate symmetry of the outward velocities indicates that takeoff is not restricted to a single direction. Bees appear to prioritize rapid dissolution before collectively flying toward the selected nest site. Bee density is highest in the *−X* half-space, which could reflect that individual fly in this direction once they have departed the swarm cluster. The order-of-magnitude reduction in trajectory counts compared to assembly reflects the much shorter timescale of disassembly.

The bee density asymmetry during both formation and dissolution suggests a net directional bias in how bees approach and depart the swarm. One possible explanation for this symmetry-breaking is that bees preferentially approach and depart from the direction of the selected nest site. During assembly, returning bees may retain memory of the swarm’s location and approach consistently from the nest-side direction. This directional bias may be further reinforced by the asymmetry in swarm shape itself. Bees on the cluster release pheromones that allow flying bees to locate the swarm from a distance [21]. A denser region of the swarm produces a stronger local signal, drawing bees preferentially from that side and creating a positive feedback loop as new arrivals further strengthen the signal. Wind can also create a global bias the transport of pheromonal cues, making it easier to locate the swarm from some directions than others [22].

## Conclusion

We developed an experimental system and 3D imaging pipeline that enables quantitative characterization of honeybee swarm assembly and disassembly dynamics in naturalistic environments. By simultaneously measuring swarm mass and morphology and reconstructing the trajectories of flying bees around the swarm, we directly observe how a cohesive cluster forms from dispersed individuals and how that structure disassembles during takeoff. In a representative swarming event, we find that bees do not approach the swarm isotropically, and instead, they move toward it along preferential directions, revealing how environmental context may shape the collective dynamics of self-assembly.

Our methods open several avenues for future investigation. Further analysis of bee trajectories could reveal whether bees are using orientation flights to create a memory of the swarm location [6]. Higher-resolution tracking of approach trajectories would allow identification of landing positions on the board or swarm surface to reveal how these initial contacts contribute to the architecture of the forming cluster. Extending tracking to bees after landing would reveal how swarm morphology emerges from the interplay of aggregation and subsequent rearrangements.

Expanding the dataset across multiple days and environmental conditions, for example wind or temperature variation, would enable comparisons of assembly and disassembly dynamics to clarify which features are robust and which vary across environmental contexts. Because a swarm makes repeated daily attempts to fly to the chosen nest site when the queen is constrained, quantifying changes in flight patterns across successive attempts may further reveal how experience, motivation, or internal state can influence the collective dynamics.

Together, these directions could provide a more complete, mechanistic picture of how thousands of individuals coordinate to build, maintain, and dissolve a living structure.

## Data availability

All code for analysis and figures is available at: https://github.com/peleg-lab/honeybee-swarm-3d-imaging. Data is available at: https://doi.org/10.5281/zenodo.18992442

## Author contributions statement

Conceptualization: D.L.C., G.N., O.P. Methodology: D.L.C., D.Z., M.K., H.R., J.S., S.H., G.N., N.R.B., O.P. Software: D.L.C. Validation: D.L.C. Formal Analysis: D.L.C. Investigation: D.L.C., D.Z., M.K., H.R., J.S., S.H., O.P. Resources: D.L.C. and O.P. Data Curation: D.L.C. Writing-Original Draft: D.L.C. and O.P. Writing-Review and Editing: D.L.C., G.N., O.P. Visualization: D.L.C. Supervision: D.L.C. and O.P. Project Administration: D.L.C. and O.P. Funding Acquisition: D.L.C. and O.P.

## Acknowledgments

We thank Hungtang Ko and Yangfan Zhang for organizing the symposium on “Animal Collective Movement in Complex Environments” at SICB 2026. We thank Ruth Williams for beekeeping support. We thank Hadley Bell Tallackson for acquiring preliminary data for this project in 2019 and Divya Pragadaraju for preliminary analysis. We acknowledge support for D.Z., H.R., and S.H. through the CU Boulder Summer Projects for Undergraduate Research and the Discovery Learning Apprenticeship Program.

## References

1. C. Anderson, G. Theraulaz, and J.-L. Deneubourg. Self-assemblages in insect societies. Insectes Sociaux, 49(2): 99–110, May 2002. ISSN 1420-9098. doi: 10.1007/s00040-002-8286-y.

2. M. Ballerini, N. Cabibbo, R. Candelier, A. Cavagna, E. Cisbani, I. Giardina, V. Lecomte, A. Orlandi, G. Parisi, A. Procaccini, M. Viale, and V. Zdravkovic. Interaction ruling animal collective behavior depends on topological rather than metric distance: Evidence from a field study. Proceedings of the National Academy of Sciences, 105(4): 1232–1237, Jan. 2008. ISSN 0027-8424, 1091-6490. doi: 10.1073/pnas.0711437105.

3. M. Beekman, R. L. Fathke, and T. D. Seeley. How does an informed minority of scouts guide a honeybee swarm as it flies to its new home? Animal Behaviour, 71(1):161–171, Jan. 2006. ISSN 00033472. doi: 10.1016/j.anbehav.2005.04.009.

4. E. Bonabeau, M. Dorigo, and G. Theraulaz. Swarm Intelligence: From Natural to Artificial Systems. Oxford University Press, 1999.

5. J. T. Bonner. A Descriptive Study of the Development of the Slime Mold Dictyostelium Discoideum. American Journal of Botany, 31(3):175–182, 1944. ISSN 1537-2197. doi: 10.1002/j.1537-2197.1944.tb08016.x.

6. E. A. Capaldi and F. C. Dyer. The role of orientation flights on homing performance in honeybees. Journal of Experimental Biology, 202(12):1655–1666, June 1999. ISSN 0022-0949, 1477-9145. doi: 10.1242/jeb.202.12.1655.

7. D. L. Chase and O. Peleg. The Physics of Sensing and Decision-Making by Animal Groups. Annual Review of Biophysics, 54(Volume 54, 2025):329–351, May 2025. ISSN 1936-122X, 1936-1238. doi: 10.1146/annurev-biophys-061824-110733.

8. J. W. Costerton, G. G. Geesey, and K.-J. Cheng. How Bacteria Stick. Scientific American, 238(1):86–95, Jan. 1978. ISSN 0036-8733. doi: 10.1038/scientificamerican0178-86.

9. I. D. Couzin. Collective cognition in animal groups. Trends in Cognitive Sciences, 13(1):36–43, Jan. 2009. ISSN 13646613. doi: 10.1016/j.tics.2008.10.002.

10. I. D. Couzin, JENS. Krause, RICHARD. James, G. D. Ruxton, and N. R. Franks. Collective Memory and Spatial Sorting in Animal Groups. Journal of Theoretical Biology, 218(1):1–11, Sept. 2002. ISSN 0022-5193. doi: 10.1006/jtbi.2002.3065.

11. I. D. Couzin, J. Krause, N. R. Franks, and S. A. Levin. Effective leadership and decision-making in animal groups on the move. Nature, 433(7025):513–516, Feb. 2005. ISSN 1476-4687. doi: 10.1038/nature03236.

12. S. M. Cully and T. D. Seeley. Self-assemblage formation in a social insect: The protective curtain of a honey bee swarm. Insectes Sociaux, 51(4):317–324, Nov. 2004. ISSN 1420-9098. doi: 10.1007/s00040-004-0743-3.

13. B. Heinrich. The Mechanisms and Energetics of Honeybee Swarm Temperature Regulation. Journal of Experimental Biology, 91(1):25–55, Apr. 1981. ISSN 0022-0949. doi: 10.1242/jeb.91.1.25.

14. G. Jocher and J. Qiu. Ultralytics YOLO11, 2024.

15. D. H. Kelley and N. T. Ouellette. Emergent dynamics of laboratory insect swarms. Scientific Reports, 3(1):1073, Jan. 2013. ISSN 2045-2322. doi: 10.1038/srep01073.

16. H. Ko, M. Hadgu, K. Komilian, and D. L. Hu. Small fire ant rafts are unstable. Physical Review Fluids, 7(9):090501, Sept. 2022. ISSN 2469-990X. doi: 10.1103/PhysRevFluids.7.090501.

17. J. Krause and G. D. Ruxton. Living in Groups. OUP Oxford, Oct. 2002. ISBN 978-0-19-850818-2.

18. M. Lindauer. Schwarmbienen auf Wohnungssuche. Zeitschrift für vergleichende Physiologie, 37(4):263–324, July 1955. ISSN 1432-1351. doi: 10.1007/BF00303153.

19. N. J. Mlot, C. A. Tovey, and D. L. Hu. Fire ants self-assemble into waterproof rafts to survive floods. Proceedings of the National Academy of Sciences, 108(19):7669–7673, May 2011. doi: 10.1073/pnas.1016658108.

20. C. Nguyen, Y. Ozkan-Aydin, H. Tuazon, D. I. Goldman, M. S. Bhamla, and O. Peleg. Emergent Collective Locomotion in an Active Polymer Model of Entangled Worm Blobs. Frontiers in Physics, 9, Sept. 2021. ISSN 2296-424X. doi: 10.3389/fphy.2021.734499.

21. D. M. T. Nguyen, M. L. Iuzzolino, A. Mankel, K. Bozek, G. J. Stephens, and O. Peleg. Flow-mediated olfactory communication in honeybee swarms. Proceedings of the National Academy of Sciences, 118(13):e2011916118, Mar. 2021. ISSN 0027-8424, 1091-6490. doi: 10.1073/pnas.2011916118.

22. D. M. T. Nguyen, G. Gharooni Fard, M. L. Iuzzolino, and O. Peleg. Gone With the Wind: Honey Bee Collective Scenting in the Presence of External Wind. In Proceedings of The ACM Collective Intelligence Conference, CI ‘23, pages 25–33, New York, NY, USA, Nov. 2023. Association for Computing Machinery. ISBN 979-8-4007-0113-9. doi: 10.1145/3582269.3615594.

23. Y. Ozkan-Aydin, D. I. Goldman, and M. S. Bhamla. Collective dynamics in entangled worm and robot blobs. Proceedings of the National Academy of Sciences, 118(6): e2010542118, Feb. 2021. doi: 10.1073/pnas.2010542118.

24. V. P. Patil, H. Tuazon, E. Kaufman, T. Chakrabortty, D. Qin, J. Dunkel, and M. S. Bhamla. Ultrafast reversible self-assembly of living tangled matter. Science, 380(6643): 392–398, Apr. 2023. doi: 10.1126/science.ade7759.

25. O. Peleg, J. M. Peters, M. K. Salcedo, and L. Mahadevan. Collective mechanical adaptation of honeybee swarms. Nature Physics, 14(12):1193–1198, Dec. 2018. ISSN 1745-2481. doi: 10.1038/s41567-018-0262-1.

26. J. M. Peters, O. Peleg, and L. Mahadevan. Thermoregulatory morphodynamics of honeybee swarm clusters. Journal of Experimental Biology, 225 (5):jeb242234, Mar. 2022. ISSN 0022-0949. doi: 10.1242/jeb.242234.225

27. M.-T. Ramsey, M. Bencsik, M. I. Newton, M. Reyes, M. Pioz, D. Crauser, N. S. Delso, and Y. Le Conte. The prediction of swarming in honeybee colonies using vibrational spectra. Scientific Reports, 10(1):9798, June 2020. ISSN 2045-2322. doi: 10.1038/s41598-020-66115-5.

28. J. Rangel and T. D. Seeley. The signals initiating the mass exodus of a honeybee swarm from its nest. Animal Behaviour, 76(6):1943–1952, Dec. 2008. ISSN 0003-3472. doi: 10.1016/j.anbehav.2008.09.004.

29. N. Ravi, V. Gabeur, Y.-T. Hu, R. Hu, C. Ryali, T. Ma, H. Khedr, R. Rädle, C. Rolland, L. Gustafson, E. Mintun, J. Pan, K. V. Alwala, N. Carion, C.-Y. Wu, R. Girshick, P. Dollár, and C. Feichtenhofer. SAM 2: Segment Anything in Images and Videos, Oct. 2024.

30. C. R. Reid, M. J. Lutz, S. Powell, A. B. Kao, I. D. Couzin, and S. Garnier. Army ants dynamically adjust living bridges in response to a cost–benefit trade-off. Proceedings of the National Academy of Sciences, 112(49):15113–15118, Dec. 2015. doi: 10.1073/pnas.1512241112.

31. R. Sarfati, J. C. Hayes, É. Sarfati, and O. Peleg. Spatio-temporal reconstruction of emergent flash synchronization in firefly swarms via stereoscopic 360-degree cameras. Journal of The Royal Society Interface, 17(170):20200179, Sept. 2020. ISSN 1742-5689. doi: 10.1098/rsif.2020.0179.

32. K. Sauer, P. Stoodley, D. M. Goeres, L. Hall-Stoodley, M. Burmølle, P. S. Stewart, and T. Bjarnsholt. The biofilm life cycle: Expanding the conceptual model of biofilm formation. Nature Reviews Microbiology, 20(10): 608–620, Oct. 2022. ISSN 1740-1534. doi: 10.1038/s41579-022-00767-0.

33. T. D. Seeley and S. C. Buhrman. Group decision making in swarms of honey bees. Behavioral Ecology and Sociobiology, 45(1):19–31, Jan. 1999. ISSN 1432-0762. doi: 10.1007/s002650050536.

34. T. D. Seeley, R. A. Morse, and P. K. Visscher. The Natural History of the Flight of Honey Bee Swarms. Psyche: A Journal of Entomology, 86(2–3):103–113, Jan. 1979. ISSN 1687-7438. doi: 10.1155/1979/80869.

35. O. Shishkov and O. Peleg. Social insects and beyond: The physics of soft, dense invertebrate aggregations. Collective Intelligence, 1(2):26339137221123758, Oct. 2022. ISSN 2633-9137. doi: 10.1177/26339137221123758.

36. O. Shishkov, C. Chen, C. A. Madonna, K. Jayaram, and O. Peleg. Strength-mass scaling law governs mass distribution inside honey bee swarms. Scientific Reports, 12(1):17388, Oct. 2022. ISSN 2045-2322. doi: 10.1038/s41598-022-21347-5.

37. M. Sinhuber, K. van der Vaart, R. Ni, J. G. Puckett, D. H. Kelley, and N. T. Ouellette. Three-dimensional time-resolved trajectories from laboratory insect swarms. Scientific Data, 6(1):190036, Mar. 2019. ISSN 2052-4463. doi: 10.1038/sdata.2019.36.

